# Mutant KRAS-associated proteome is mainly controlled by exogenous factors

**DOI:** 10.1101/2022.01.05.475056

**Authors:** Patrícia Dias Carvalho, Flávia Martins, Joana Carvalho, Maria José Oliveira, Sérgia Velho

## Abstract

KRAS signaling has been extensively studied, yet the clarification between KRAS-autonomous and non-autonomous mechanisms are still less explored. Understanding how KRAS signaling and effects are affected by exogenous stimuli can provide valuable insights not only to understand resistance mechanisms that justify pathway inhibition failure, but also to uncover novel therapeutic targets for mutant KRAS patients. Hence, aiming at understanding KRAS-autonomous versus non autonomous mechanisms, we studied the response of two mutant KRAS colorectal cancer cell lines (HCT116 and LS174T) - control and KRAS silenced- to TGFβ1-activated fibroblasts secretome. By performing a total proteome analysis, we observed that TGFβ1-activated fibroblast-secreted factors triggered cell line-specific proteome alterations and that mutant KRAS governs approximately 1/3 of those alterations. Moreover, the analysis of the impact of exogenous factors on the modulation of KRAS proteome revealed that, in both cell lines, more than 2/3 of the KRAS-associated proteome is controlled in a KRAS-non-autonomous manner and dependent on the exogenous factors. This work highlights the context-dependency of KRAS-associated signaling and reinforces the importance of establishing more integrative models resembling the complexity of the tumor microenvironment to study KRAS-associated signals.

## 1. Introduction

KRAS belongs to the RAS family of small GTPases. Owing to its inner membrane location, KRAS functions as a transducer of extracellular stimuli to the interior of the cell. By integrating signals from different tyrosine kinase receptors (RTKs), KRAS governs several distinct signaling cascades, such as RAF–MEK–ERK, PI3K–AKT–mTOR, and RALGDS–RAL, dictating cell fate. Missense substitutions occurring in KRAS gene, either reduce GTP hydrolysis or increase the rate of GTP loading, altering its ON/OFF homeostasis towards the active state, thus driving cell proliferation and survival [1,2].

Standing as the most frequently mutated oncogene in human cancer, with particular relevance in pancreatic, colorectal (CRC) and lung cancers [2], KRAS is considered a key therapeutic target. However, despite of extensive attempts to target KRAS or its downstream signaling effectors, only the recently approved KRAS G12C specific inhibitor demonstrated clinical efficiency [3]. So, one must interrogate why so many different drugs and strategies failed to show therapeutic efficiency when reaching the clinical setting. Despite the alterations induced by specific mutations, mutant KRAS (mutKRAS) forms still depend, to a certain extent, on the activation by external stimuli [4]. Moreover, different mutations display distinct effector affinities and levels of activation, dictating diverse effects [2,5–7]. Likewise, KRAS downstream effects have been shown to be context-dependent, showing allele and tissue/tumor specificities [8,9]. In addition, it has been demonstrated that signaling heterogeneity in both the tumor and microenvironment impacts treatment response [10]. Therefore, the study of the crosstalk of mutKRAS cancer cells with the tumor microenvironment (TME) components acquires a special relevance. In fact, mutKRAS cancer cells have been shown to communicate with and modulate the TME, favoring tumor progression and malignancy [11–15]. As so, in vitro studies evaluating drug responses only considering cancer cells in their optimal culture conditions, are very reductionist and might not generate translational knowledge. Thus, more integrative studies are needed to better understand the impact of the microenvironment on KRAS-driven signaling and therapy response/resistance. In line with this, our group has recently shown that mutKRAS can regulate functional effects autonomously or it can cooperate with fibroblast-secreted factors to modulate cancer cell invasive behavior [16].

Herein, we aimed to demonstrate the impact of microenvironmental cues on KRAS-driven signaling. As fibroblasts are one of the major components of the TME, we used TGFβ1-activated fibroblast-derived secretome as a source of microenvironment signals. We demonstrated that mutKRAS controls both autonomous, and non-autonomous fibroblast-dependent signaling. Noteworthy, we showed that 2/3 of the total proteome regulated by mutKRAS are stimuli-dependent. Moreover, fibroblast-derived signals even reversed the expression trend of some KRAS-regulated proteins when comparing stimulated with non-stimulated cells. Overall, our data shows that the mutKRAS-associated proteome profile drastically changes in response to external stimulation, suggesting that its oncogenic signaling is mainly regulated in a non-autonomous manner. As such, we propose that studies addressing the oncogenic effects of KRAS, the identification of therapeutic targets or biomarkers of therapy resistance should take into consideration the influence of the microenvironment in dictating KRAS signaling.

## 2. Materials and Methods

### Cell culture

HCT116 colorectal cancer cell line and CCD-18Co normal colon fibroblasts were purchased from American Type Culture Collection (ATCC). LS174T cells were kindly provided by Dr. Ragnhild A. Lothe (Oslo University Hospital). HCT116 cells were cultured in RPMI 1640 medium (Gibco, Thermo Fisher Scientific) and LS174T and CCD-18Co were cultured in DMEM medium (Gibco, Thermo Fisher Scientific). For all cell lines the respective medium was supplemented with 10% fetal bovine serum (Hyclone) and 1% penicillin–streptomycin (Gibco, Thermo Fisher Scientific). Cells were maintained at 37°C in a humidified atmosphere with 5% CO2.

### CCD-18Co conditioned media production

For conditioned media production, the same number of cells was plated in two T75 culture flasks and cultured until approximately 90% of confluence. At the desired confluence, cells were washed twice with PBS buffer and new media was added. “Normal-like” fibroblasts were cultured in serum-free DMEM (supplemented only with 1% penicillin/streptomycin) and “activated-fibroblasts” were cultured in the same medium supplemented with 10 ng/mL of rhTGFβ1 (ImmunoTools GmbH). As control media, DMEM+1% penicillin/streptomycin and DMEM+1% penicillin/streptomycin+ 10 ng/mL rhTGF-β1-were added to culture flasks without cells. After four days in optimal culture conditions, conditioned media (CM) were harvested, centrifuged, filtered through a 0,2 μm filter and stored at −20°C until use. Cells were tripsinized and counted to assure an equivalent number of cells in both conditions. Total protein was extracted, and fibroblast-activation was confirmed through the evaluation of alpha smooth muscle actin (α-SMA) expression by western blot (Figure S1).

### Gene silencing by siRNA transfection and treatment with conditioned media

Cells were seeded in six-well plates (150,000 and 200,000 cells for HCT116 and LS174T, respectively) and transfected after approximately 16h, using Lipofectamine RNAiMAX (Invitrogen, Thermo Fisher Scientific) in reduced-serum Opti-MEM medium (Gibco, Thermo Fisher Scientific), following manufacturer’s instructions. Gene silencing was achieved with ON-TARGETplus SMARTpool small interfering RNA specific for KRAS (L-005069-00-0010, Dharmacon) at a final concentration of 10 nM. A non-targeting siRNA (D-001810-01-50; ON-TARGETplus Non-targeting siRNA #1, Dharmacon) was used as a negative control. 72h after transfection, control (siCTRL) and KRAS silenced (siKRAS) HCT116 and LS174T cells, were washed and treated with serum free CM from activated fibroblasts and the respective control, during 24h. KRAS silencing efficiency was monitored by western blot (Figure S2). For each cell line, three independent biological replicates were performed.

### Protein extraction and western blotting

Total protein was extracted using ice cold RIPA Buffer [25mM Tris-HCl pH=7-8; 150mM NaCl; 0.5% sodium deoxycholate; 1% triton X-100] supplemented with a pro-tease inhibitor cocktail (Roche) and a phosphatase inhibitor cocktail (Sigma-Aldrich). Protein concentration was determined using the DCProtein assay kit from BioRad and 100µg of total protein were processed for proteomics analysis. Furthermore, 25µg of protein were resolved on sodium dodecyl sulphate-polyacrylamide gel electrophoresis (SDS-PAGE) under denaturing conditions and transferred to Protran Premium NC 0.45µm membranes (Amersham Biosciences, GE Healthcare). Membranes were blocked for 1h at RT and incubated overnight at 4°C with agitation, with the respective primary antibody against KRAS (LS-Bio, LS-C175665; 1:4000), α-SMA (Abcam, ab7817, 1:250) or GAPDH (Santa Cruz Biotechnology, sc-47724, 1:10000), all diluted in 5% non-fat milk in PBS+ 0.5% Tween 20. After incubation with the specific anti-mouse (NA931, GE Healthcare) HRP-conjugated secondary antibodies for 1h at RT, bands were detected using ECL (BioRad) and film sheets exposure (Amersham Biosciences, GE Healthcare).

### Sample processing for proteomic analysis

Sample processing and proteomic analysis was performed at i3S Proteomics Scientific Platform. For proteomic analysis, a single-step reduction and alkylation with tris-2(-carboxyethyl)-phosphine (TCEP)/ chloroacetamide (CAA) was performed in combination with the single-pot solid-phase-enhanced sample preparation (SP3) protocol as described elsewhere [17,18]. Briefly, to reduce disulfide bonds and alkylate cysteines, sodium deoxycholate (SDC) 2x buffer (200 mM Tris pH 8.5, SDC 2%, 20 mM TCEP, 80 mM CAA) was added to the protein sample and incubated for 10 min at 95°C, 1000 rpm. Next, 100 µL of Sera-Mag Magnetic Beads (10 µg/µL, GE Healthcare) and ethanol 100% were added to the protein solution (50% ethanol, final concentration), resuspended, and incubated at 22°C for 10 min at 1000 rpm. After complete binding, the samples were washed with 80% ethanol. To achieve protein enzymatic digestion, 50 µL of triethylammoniumbicarbonate (TEAB) 50 mM mixed with Trypsin + LysC (2 µg) was added to the beads and incubated overnight at 37°C with 1000 rpm agitation. In the next day, 1.3 mL of 100% acetonitrile (ACN) was added to samples and incubated for 20 min at 22ºC, 1000 rpm. Tubes were placed in a magnetic rack and beads were washed with of 100% ACN. Following, 100 µL of TEAB 50 mM was added to the beads, resuspended, and incubated for 5 min at 22°C, 1000rpm. After incubation, tubes were placed in magnetic rack until the beads have migrated to the tube wall and the supernatant was transferred to a new tube, with 20 µL of 5% formic acid (FA). Afterwards, tubes with FA were placed in a SpeedVac until the samples were dry. Then, samples were resuspended in 0.1% FA and peptides were quantified by Pierce™ Quantitative Fluorometric Peptide Assay (Thermo Fisher Scientific). 500 ng of peptides of each sample were analyzed by Liquid Chromatography-Mass Spectrometry (LC-MS).

### LC-MS/MS-analysis

Protein identification and quantitation was performed by nanoLC–MS/MS using an Ultimate 3000 liquid chromatography system coupled to a Q-Exactive Hybrid Quadrupole-Orbitrap mass spectrometer (Thermo Fisher Scientific), following published protocols [19]. Specifically, samples were loaded onto a trapping cartridge (Acclaim PepMap C18 100Å, 5 mm x 300 µm i.d., 160454, Thermo Fisher Scientific) in a mobile phase of 2% ACN, 0.1% FA at 10 µL/min. After 3 min loading, the trap column was switched in-line to a 50 cm by 75μm inner diameter EASY-Spray column (ES803, PepMap RSLC, C18, 2 μm, Thermo Fisher Scientific) at 300 nL/min. Separation was generated by mixing A: 0.1% FA, and B: 80% ACN, with the following gradient: 5 min (2.5% B to 10% B), 120 min (10% B to 30% B), 35 min (30% B to 50% B), 3 min (50% B to 99% B) and 12 min (hold 99% B). Data acquisition was controlled by Xcalibur 4.0 and Tune 2.9 software (Thermo Fisher Scientific).

The mass spectrometer was operated in data-dependent (dd) positive acquisition mode alternating between a full scan (m/z 380-1580) and subsequent HCD MS/MS of the 10 most intense peaks from full scan (normalized collision energy of 27%). ESI spray voltage was 1.9 kV. The following settings were used: global settings-use lock masses best (m/z 445.12003), lock mass injection Full MS, chrom. peak width (FWHM) 15s; full scan settings-70k resolution (m/z 200), AGC target 3e6, maximum injection time 120 ms; dd settings-minimum AGC target 8e3, intensity threshold 7.3e4, charge exclusion: unassigned, 1, 8, >8, peptide match preferred, exclude isotopes on, dynamic exclusion 45s; MS2 settings-microscans 1, resolution 35k (m/z 200), AGC target 2e5, maximum injection time 110 ms, isolation window 2.0 m/z, isolation offset 0.0 m/z, spectrum data type profile.

### Data and Bioinformatics Analysis

The acquired raw data were analyzed using the Proteome Discoverer 2.5.0.400 software (Thermo Scientific) and searched against the UniProt database for the Homo sapiens Proteome 2020_05 (75069 entries). The Sequest HT search engine was used to identify tryptic peptides. The ion mass tolerance was 10 ppm for precursor ions and 0.02 Da for fragment ions. Maximum allowed missing cleavage sites was set 2. Cysteine carbamidomethylation was defined as constant modification. Methionine oxidation, serine, threonine and tyrosine phosphorylation and protein N-terminus acetylation were defined as variable modifications. Peptide confidence was set to high. The processing node Percolator was enabled with the following settings: maximum delta Cn 0.05; decoy database search target FDR 1%, validation based on q-value. Protein label free quantitation was performed with the Minora feature detector node at the processing step. Precursor ions quantification was performing at the processing step with the following parameters: unique plus razor peptides were considered for quantification, precursor abundance was based on intensity, normalization mode was based on total peptide amount, protein ratio calculation was pairwise ratio based, imputation was not performed, hypothesis test was based on t-test (background based). The mass spectrometry proteomics data have been deposited to the ProteomeXchange Consortium via the PRIDE [17] partner repository with the dataset identifier PXD030551 and 10.6019/PXD030551. Gene Ontology (GO) analysis and Kyoto Encyclopedia of Genes and Genomes (KEEG) pathway analysis was performed using DAVID Bioinformatics. For these analysis, p-value cutoff of 0.05 was applied. Graphics were made using GraphPad Prism version 9.0.0 and the enrichment score was calculated by –log P-value.

## 3. Results

To investigate the impact of fibroblast-derived microenvironmental factors on KRAS-driven signaling, we profiled the total proteome of two distinct mutKRAS CRC cell lines (HCT116 and LS174T) upon KRAS silencing, followed by treatment with conditioned medium of rhTGFβ1-activated fibroblasts (siKRAS_FibCM), mimicking CAFs secretome. As controls, we used non-targeting (siCTRL) and KRAS silenced (siKRAS) cells cultured with control CM (siCTRL_ctrlCM; siKRAS_ctrlCM) as well as siCTRL cells cultured with FibCM (siCTRL_FibCM).

Principal component (PC) analysis revealed striking differences between the two cell lines. While all conditions belonging to the same cell line cluster together, the two different cell lines can be discriminated by PC 2, with 23.8% of the observed differences (Figure 1a). Analyzing the cell lines individually, PCA plots revealed that it was possible to discriminate between siCTRL and siKRAS in HCT116 cells by PC 2, with 13.6% of the observed differences, although the same was not possible to distinguish between cells treated with ctrlCM or FibCM (Figure 1b). Regarding LS174T cells, it was not possible to completely discriminate between different conditions, although most of siCTRL cells were separated from siKRAS group (Figure 1c).

**Figure 1:**
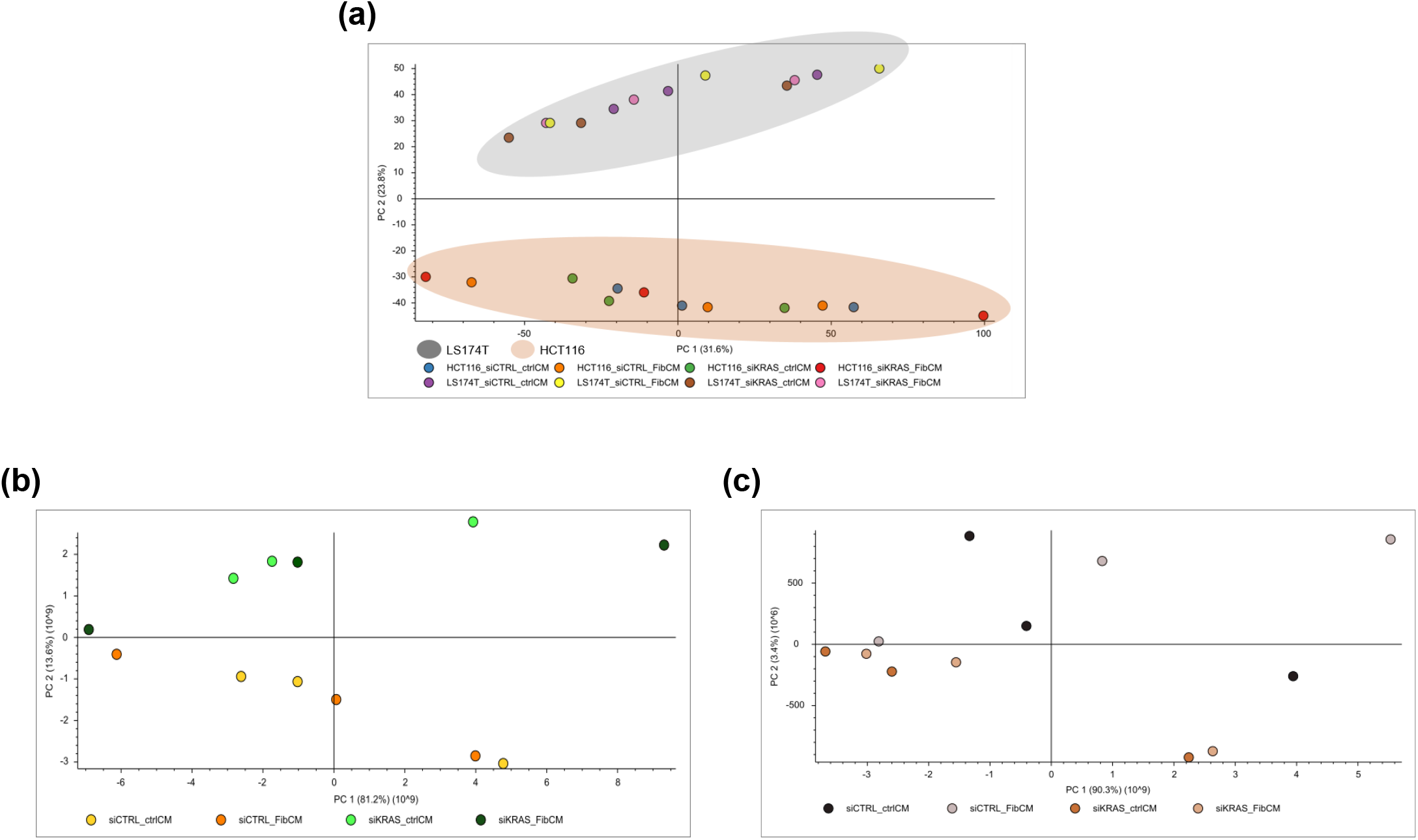
Principal component analysis (PCA) of total identified proteins in HCT116 and LS174T cell lines. (a) PCA plot of the two cell lines together (b) PCA plot for HCT116 cell line (c) PCA plot for LS174T cell line. siCTRL-non-targeting siRNA control cells; siKRAS-KRAS silenced cells; ctrlCM DMEM+ TGFβ1 control medium: FibCM-TGFβ1-activated fibroblasts conditioned medium.

Because cancer-associated fibroblasts are well-known modulators of cancer cell malignant traits, we performed a detailed in silico functional characterization of the differentially expressed proteins by means of Gene Ontology (GO) and KEGG pathway enrichment in order to determine: (i) the overall influence of fibroblast-secreted factors on the proteome of CRC cell lines; (ii) to which extent these alterations depend on oncogenic KRAS; and (iii) the effect of the fibroblast-secreted factors in conditioning oncogenic KRAS signaling, specifically. The workflow of the performed analysis is depicted in Figure 2.

**Figure 2:**
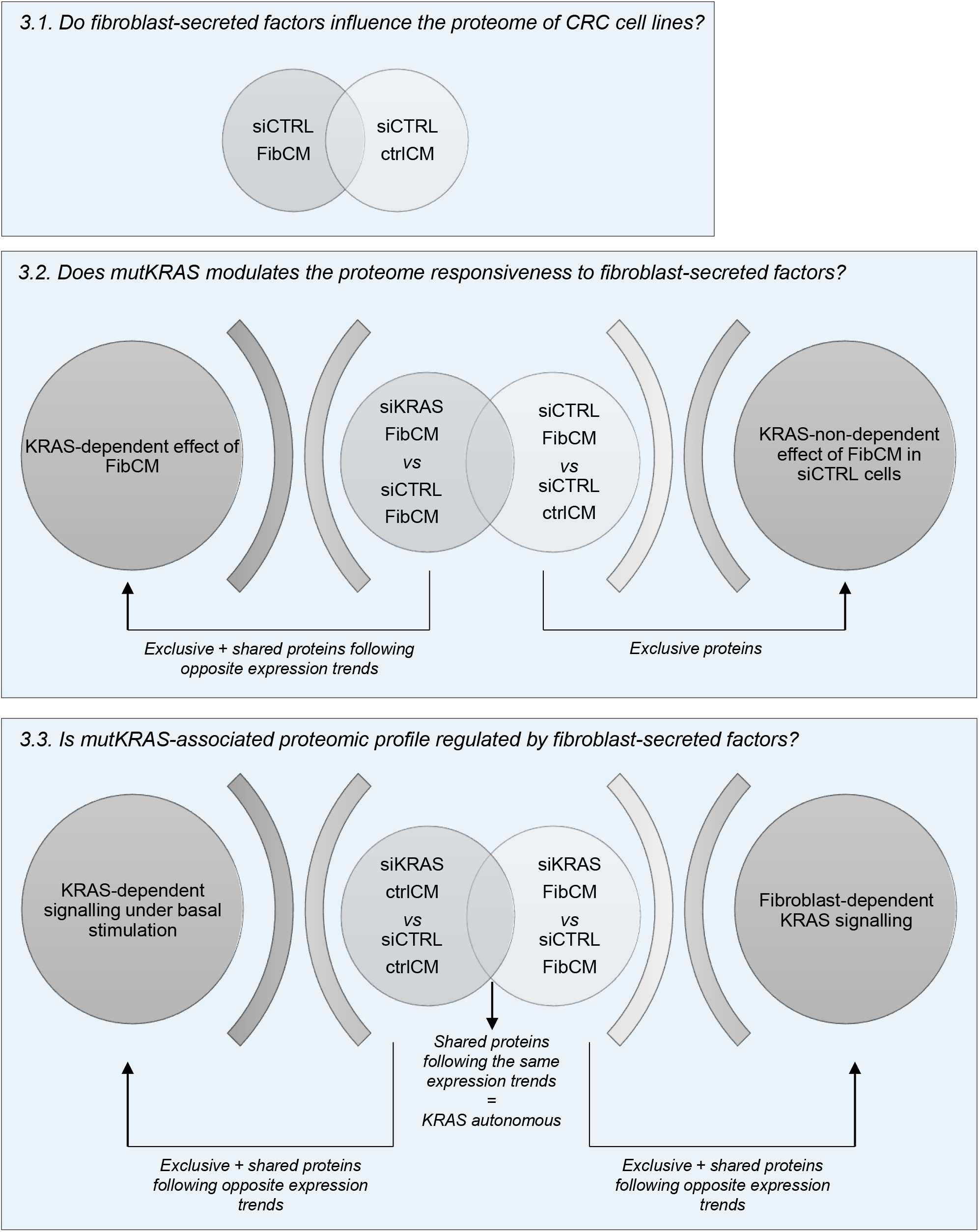
Schematic representation of the workflow analysis.

### 3.1. Fibroblast-secreted factors impact the proteome of CRC cell lines

First, we analyzed the impact of TGFβ1-activated fibroblast-derived factors on the protein expression profile of HCT116 and LS174T CRC cell lines. To do so, we compared differentially expressed proteins between siCTRL cells cultured with FibCM or ctrlCM. Considering the effect of FibCM in HCT116 cells, 77 proteins were found to be differentially expressed, with 42 upregulated and 35 downregulated (Figure 3a and Supplementary File S1); in LS174T cells, 69 proteins were found to be differentially expressed, with 35 upregulated and 34 downregulated (Figure 3b and Supplementary File S2).

**Figure 3:**
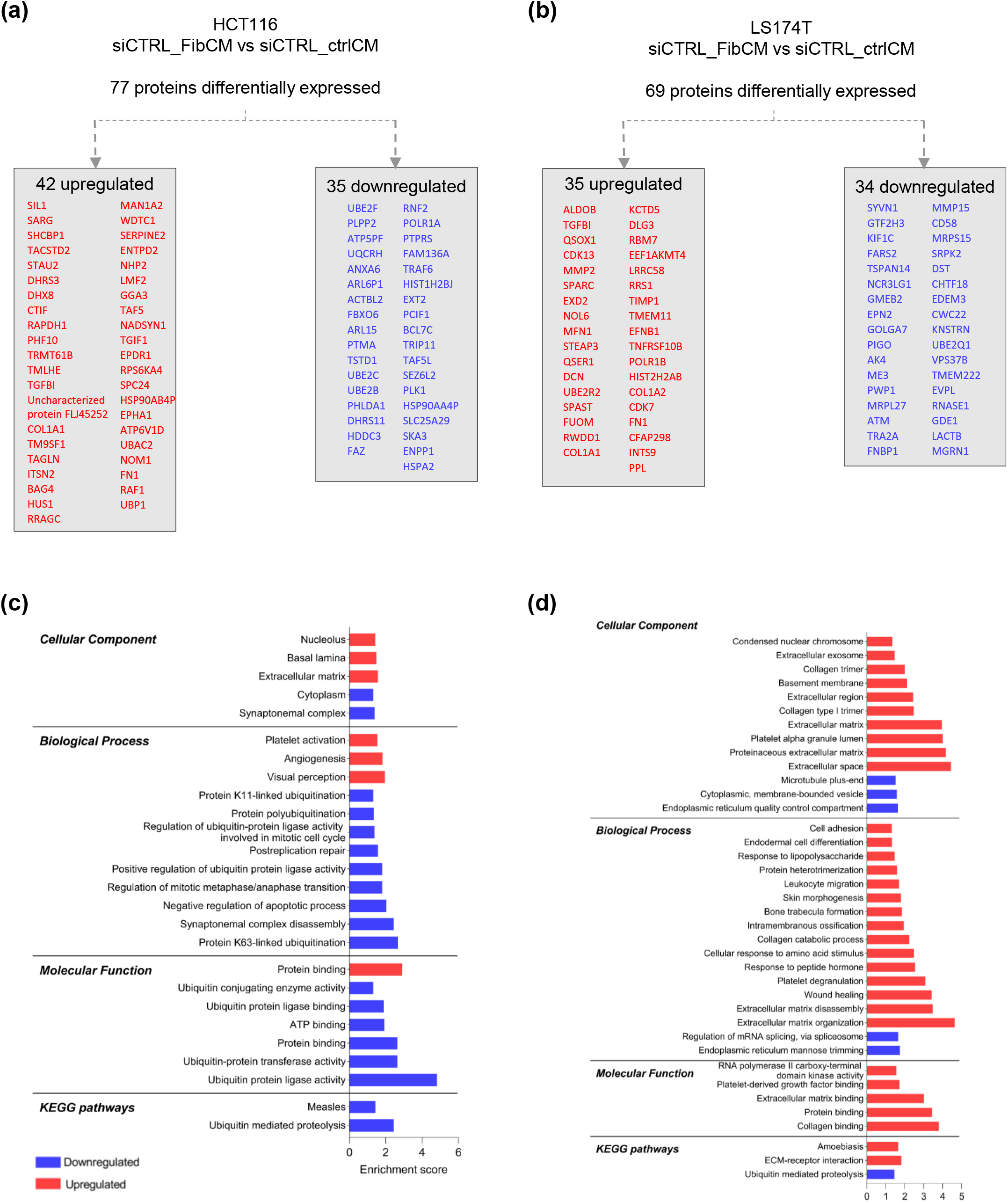
HCT116 and LS174T differentially expressed proteins, upon culture with TGFβ1-activated fibroblasts conditioned medium. Scheme showing significantly up and downregulated proteins in HCT116 (a) and LS174T (b) when comparing cells cultured in ctrlCM and FibCM. GO and KEGG pathways analysis of up and downregulated proteins in HCT116 (c) and LS174T (d) when comparing cells cultured in ctrlCM and FibCM. The enrichment score was calculated by –log P-value.

GO analysis revealed that proteins associated with the Extracellular Matrix (ECM) were significantly upregulated in both cell lines, upon culture with FibCM (Figure 3c and 3d). Of note, only three differentially expressed proteins were found to be commonly regulated by FibCM in both cell lines and belong to the ECM category: Collagen Type I Alpha 1 (COL1A1), Fibronectin (FN1), and Transforming Growth Factor Beta Induced (TGFBI). Further, HCT116 cells also presented an upregulation of proteins localized at the nucleolus and at the basal lamina (Figure 3c). In LS174T, proteins upregulated in response to FibCM were also localized at the extracellular space, at the basement membrane, at the mitochondria, and other cellular components (Figure 3c). Downregulated proteins in HCT116 cells were associated with synaptonemal complex and cytoplasm localizations, while for LS174T those were related to the endoplasmic reticulum quality control compartment, the cytoplasmic membrane-bounded vesicles, and the microtubule plus-end (Figure 3c and Figure 3d). Concerning the biological processes, whilst upregulated proteins of HCT116 cells were mainly involved in angiogenesis; in LS174T cells, upregulated proteins were spanned across several processes, such as ECM disassembly/organization, wound healing, collagen catabolic process and cell adhesion. The downregulated proteins were involved in several ubiquitination processes, negative regulation of apoptosis, and processes involved in cell division in HCT116 cells, while in LS174T cells these proteins were involved in mRNA splicing regulation and endoplasmic reticulum mannose trimming (Figure 3c and Figure 3d). Regarding the molecular function, in HCT116 cells, some proteins involved in protein binding were found upregulated while others were downregulated; proteins mainly involved in processes impacting ubiquitin were downregulated (Figure 3c). As for LS174T cells, only upregulated proteins were significantly involved in collagen, protein, extracellular matrix and platelet-derived growth factor binding, and RNA polymerase II carboxy-terminal domain kinase activity (Figure 3d). KEGG pathway analysis revealed that solely upregulated pathways presented statistically significant results for LS174T cells, including the relevant ECM-receptor interactions. The downregulated proteins were mainly involved in ubiquitin mediated proteolysis pathways in both cell lines (Figure 3c and Figure 3d). Together this data shows that TGFβ1-activated fibroblasts secretome alters CRC cells proteome, mainly in a cell line-dependent manner.

### 3.2. MutKRAS modulates the proteomic profile associated with cancer cell response to fibroblast-secreted factors

Next, we evaluated the influence of mutKRAS on the modulation of the proteome responsiveness to fibroblast-secreted factors. To do so, we first determined the impact of FibCM in the proteomic profile of siKRAS cells by comparing siKRAS vs siCTRL cultured in FibCM, and then compared it to the proteomic profile previously obtained from the siCTRL_FibCM vs siCTRL_ctrlCM analysis. From the 77 proteins that were differentially expressed by HCT116 siCTRL cells in response to fibroblast-secreted factors, 33 (43%) were also found to be altered in siKRAS cells cultured with FibCM (Figure 4a). Most of these common proteins, 26/33 (79%) were controlled by KRAS as their expression followed opposite trends in siKRAS cells compared to the siCTRL (Figure 4b); 7/33 (21%) experienced an increment on their expression tendency upon KRAS silencing, suggesting that KRAS antagonizes their expression in response to fibroblast-secreted factors (Figure 4c). Overall, from the 77 proteins differently expressed upon treatment with FibCM, 44 (57%) were independent of KRAS, 26 (34%) were positively controlled by KRAS, and 7 (9%) were antagonized by KRAS (Figure 4a-d). GO terms analysis of the 26 proteins controlled by KRAS revealed that upregulated proteins in siKRAS cells, were associated with the synaptomenal complex, spindle microtubule and cell surface localization, whereas downregulated proteins were mainly localized at the nucleolus and nucleoplasm. For the biological processes, only upregulated proteins were significantly associated with synaptomenal complex disassembly and response to unfolded protein. At the molecular function level, upregulated proteins were related to protein binding, and downregulated proteins with poly(A) RNA binding (Figure 4b). GO terms analysis of the seven proteins with the same tendency, revealed their presence at extracellular exosomes (Figure 4c), thus suggesting that fibroblast-secretome can be prompting cells to increase their intercellular communication via exosomes. Twenty-four out of the 44 proteins of siCTRL cells exclusively identified as an effect of the FibCM, were upregulated and found to be related with ECM localization, biological processes of cellular response to epidermal growth factor stimulus and substrate adhesion-dependent cell spreading; to protein binding at the molecular function level and to focal adhesion in the KEGG pathway analysis (Figure 4d). The downregulated proteins (20/44), were localized at the mitochondria, related with protein-ubiquitination processes, and associated to ubiquitin-related activities. In accordance, KEGG pathway analysis showed that these proteins were found in ubiquitin mediated proteolysis pathway (Figure 4d).

**Figure 4:**
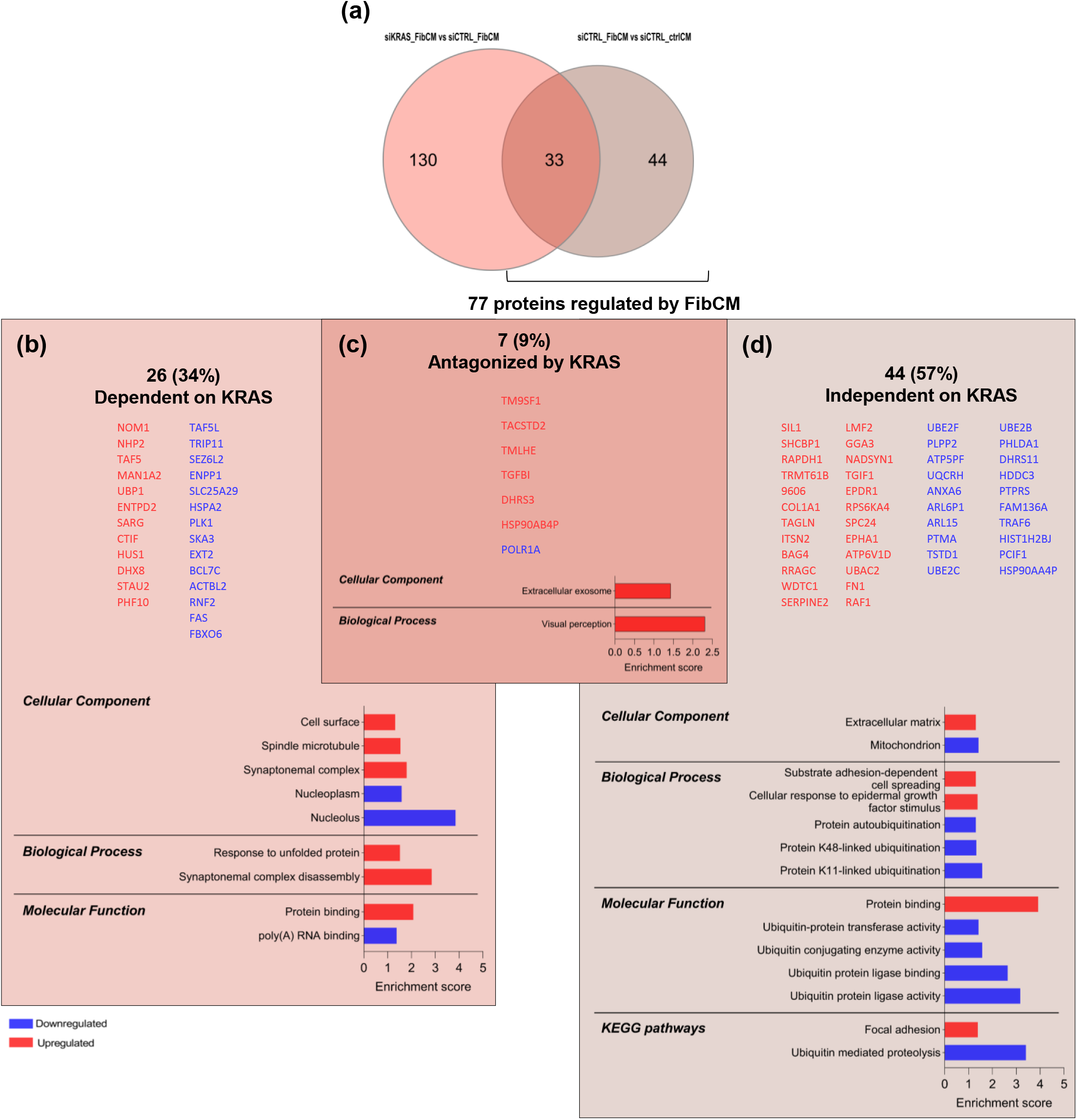
Analysis of differentially expressed proteins in HCT116 cells, when comparing siKRAS_FibCM vs siCTRL_FibCM and siCTRL_FibCM vs siCTRL_ctrlCM. (a) Venn diagram showing exclusive and common proteins identified in the analyzed conditions. (b-d) GO and KEGG pathways analysis of: common proteins following opposite trends (b), common proteins following the same tendency (c) and proteins exclusively found as an effect of the FibCM in siCTRL cells (d). The enrichment score was calculated by –log P-value.

From the 69 proteins found differentially expressed in LS174T siCTRL cells in response to fibroblast-secreted factors, 26 (37%) were also found to be altered in siKRAS cells cultured with FibCM (Figure 5a). From these, three (11%) followed the same tendency and 23 (88%) followed opposite tendencies, thus highlighting the important role of mutKRAS in regulating the response to external factors. Overall, from the 69 proteins differently expressed upon treatment with FibCM, 43 (62.4%) are independent of KRAS, 23 (33.3%) are positively controlled by KRAS, and three (4.3%) are antagonized by KRAS (Figure 5a-d). GO terms analysis demonstrated that, only the upregulated proteins from the 23 controlled by KRAS, displayed significative results at the cellular component and biological process levels, being related to extracellular exosome localization and endocytosis process (Figure 5b). From the 43 proteins of siCTRL cells exclusively identified upon culture with FibCM, the 24 upregulated were mainly localized at the extracellular space, extracellular exosome, ECM and extracellular region among other localizations while the downregulated proteins (19/43) were intracellular, being located at the mitochondrial matrix and the endoplasmic reticulum quality control compartment (Figure 5d). As for the biological processes, upregulated proteins were shown to be involved in a multitude of processes, such as extracellular matrix organization and disassembly, cell adhesion, wound healing, angiogenesis and blood vessel development, collagen catabolic process and fibril organization among others, whereas downregulated proteins only displayed two terms: endoplasmic reticulum mannose trimming and endoplasmic reticulum unfolded protein response (Figure 5d). Concerning the molecular function, upregulated proteins were involved in protein, collagen, platelet-derived growth factor and ECM binding, while the downregulated ones were involved in calcium ion binding. KEGG pathways analysis showed that only upregulated proteins were significantly involved in ECM-receptor interaction, proteoglycans in cancer and focal adhesion (Figure 5d).

**Figure 5:**
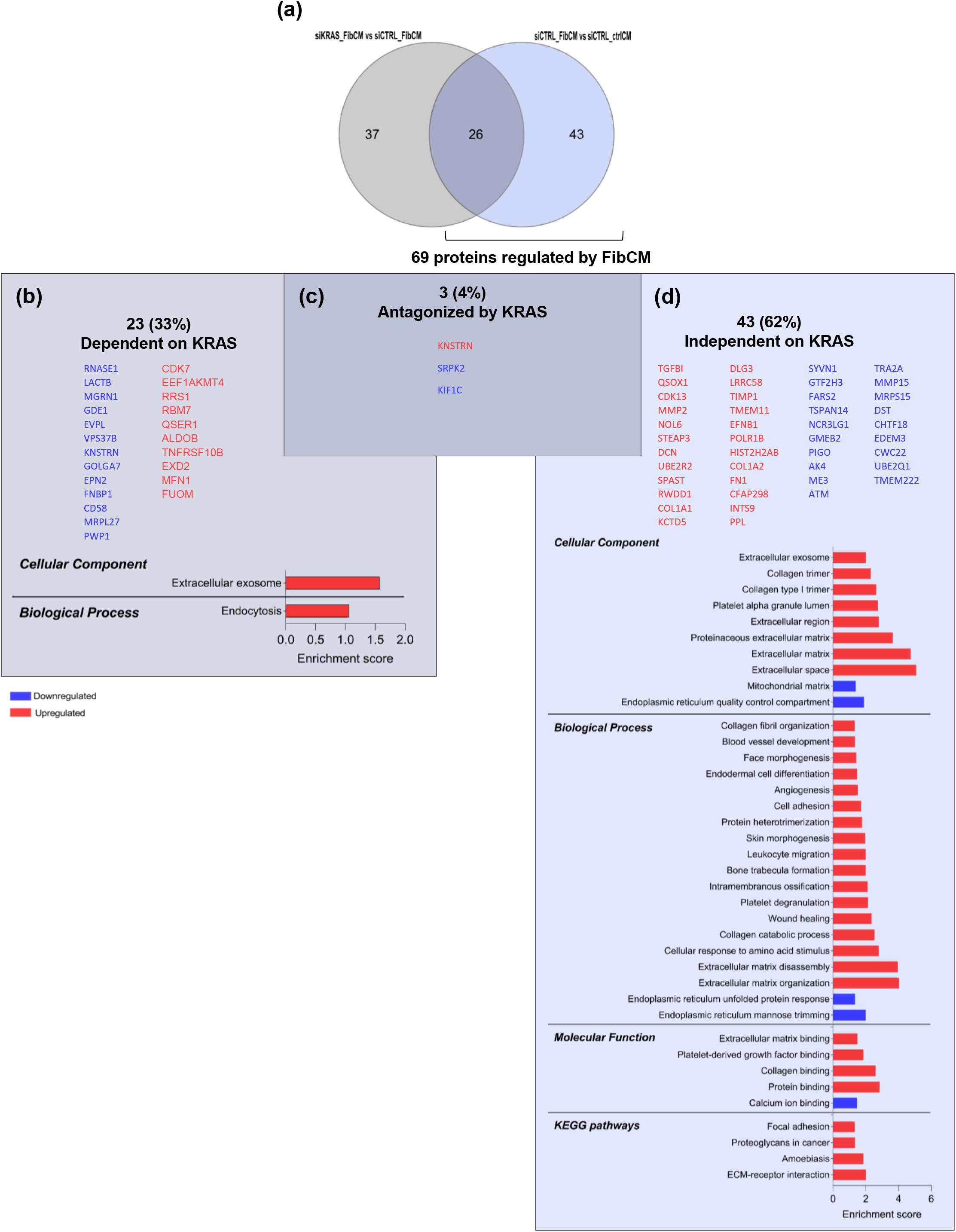
Analysis of differentially expressed proteins in LS174T cells, when comparing siKRAS_FibCM vs siCTRL_FibCM and siCTRL_FibCM vs siCTRL_ctrlCM. (a) Venn diagram showing exclusive and common proteins identified in the analyzed conditions. (b-d) GO and KEGG pathways analysis of: common proteins following opposite trends (b) common proteins following the same tendency, showing no significative results (c), and proteins exclusively found as an effect of the FibCM in siCTRL cells (d). The enrichment score was calculated by –log P-value.

Gathering these results, we can understand that within control cells, the proteome that is modulated by the fibroblasts-secreted factors, and is dependent on KRAS signaling, correspond to 43% and 37,7% of the differentially expressed proteins identified in HCT116 and LS174T cell lines, respectively. Moreover, many of the changes induced by fibroblast-secreted factors that are KRAS-independent, are related to cell-ECM interactions.

### 3.3. The proteomic profile associated with mutKRAS is highly regulated by fibroblast-secreted factors

As fibroblast-secreted factors modulated the proteome of CRC cells, in general, we next questioned whether those factors might also underly specific proteomic responses associated with oncogenic KRAS. Therefore, we compared the proteome of: (i) siKRAS vs siCRTL cells cultured in ctrlCM, unveiling KRAS-dependent signalling under basal stimulation; and (ii) siKRAS vs siCTRL cells cultured in FibCM, to dissect a fibroblast-dependent mutKRAS signalling. A comparative analysis of these two datasets will pinpoint KRAS-non autonomous changes, as alterations in mutKRAS signaling will result from stimulation with fibroblast-secretome.

In HCT116 cell line cultured with ctrlCM, KRAS silencing significantly altered the expression of 172 proteins (70 upregulated and 102 downregulated) comparing to siCTRL cells. The comparison between siKRAS vs siCTRL proteome upon culture with FibCM resulted in 163 differently expressed proteins (88 upregulated and 75 downregulated) (Figure 6a and Supplementary File S1). Moreover, a comparative analysis of these two datasets highlighted 109 proteins exclusively found in siKRAS cells cultured in ctrlCM (109/172 – 63%), 100 proteins exclusively of siKRAS cells cultured in FibCM (100/163 – 61%), and 63 proteins that were shared by the two conditions (Figure 6a). Fifty-four of the 63 shared proteins (86%) followed the same expression tendency (either up or downregulation), and nine (14%) showed opposite expression tendencies (Figure 6a-c and Supplementary File S1). Hence, these nine proteins were also included in the panel of exclusive proteins, despite being common to both culture conditions. As so, in HCT116 cell line, fibroblast-secreted factors control 67% (100 +9/163) of the proteome associated to oncogenic KRAS. Six out of the nine proteins (SARG, CTIF, PHF10, BLOC1S5, SPINDOC and HUS1) with opposite expression were upregulated upon KRAS silencing in ctrlCM and became downregulated upon culture of siKRAS cells with FibCM. The remaining three (PLK1, SKA3, RTN2) were downregulated upon KRAS silencing in ctrlCM and became upregulated upon culture with FibCM. Upregulated proteins were related to spindle microtubule and kinetochore localizations and were involved in mitotic nuclear division (Figure 6b). The 54 shared proteins are likely resulting from KRAS-autonomous signaling, since their up or downregulation tendencies were maintained independently of the culture conditions. In silico functional analysis of these shared proteins revealed that only the upregulated ones were significantly associated with the following cellular components: cytosol, extracellular exosomes, extracellular matrix and cytoplasm. Considering the relevant biological processes in this context, the upregulated proteins were involved in retinoic acid/retinol biosynthesis and metabolism, and type I interferon pathway, while the downregulated proteins were involved in aminoacid biosynthesis, response to glycose starvation, and protein kinase B signaling. Accordingly, the molecular function category highlighted NADP-retinol and retinol dehydrogenase activity in upregulated proteins, while downregulated proteins were involved in identical protein binding. Moreover, KEGG pathway analysis revealed that upregulated proteins were involved in metabolic pathways while the downregulated ones were involved in alanine, aspartate and glutamate metabolism (Figure 6c). Noteworthy, the vast majority of the total proteome associated to oncogenic KRAS was found to be highly dependent on external stimulation (109/163 – 67% induced by FibCM, and 118/172 – 69% induced by CtrlCM), resulting in differential regulation of several biologic processes through a KRAS-dependent-non-autonomous signaling.

**Figure 6:**
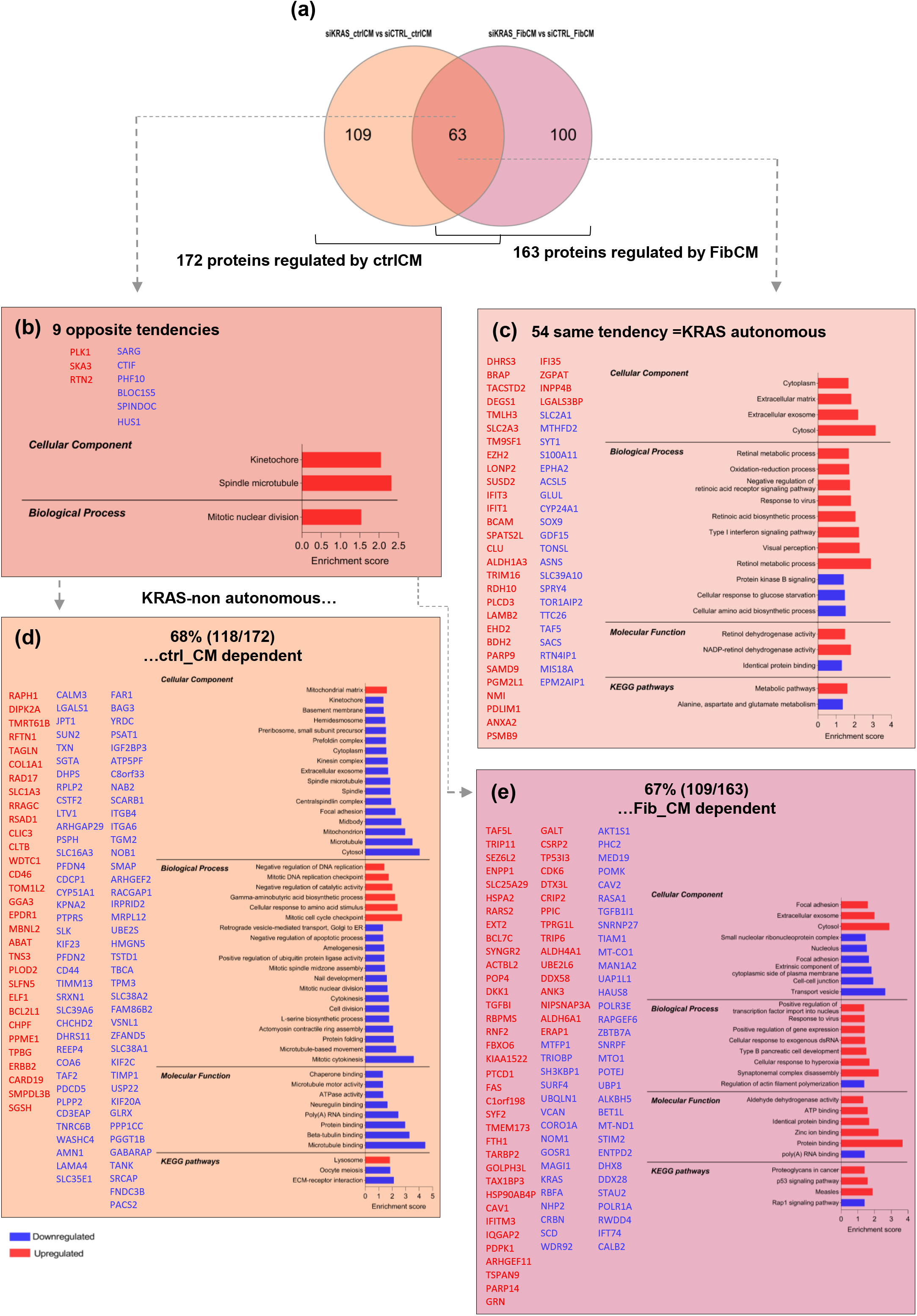
Analysis of differentially expressed proteins in HCT116 cells, when comparing siKRAS_ctrlCM vs siCTRL_ctrlCM and siKRAS_FibCM vs siCTRL_FibCM. (a) Venn diagram showing exclusive and common proteins identified in the analyzed conditions. (b-e) GO and KEGG pathways analysis of: common proteins following opposite trends (b), common proteins following the same tendency (c), exclusive proteins together with the commons that followed opposite tendencies found in the comparison of siKRAS vs siCTRL cultured in ctrlCM (d) and exclusive proteins together with the commons that followed opposite tendencies found in the comparison of siKRAS vs siCTRL cultured in FibCM (e). The enrichment score was calculated by –log P-value.

In particular, a detailed analysis of the 109 exclusive proteins of siKRAS_ctrlCM cells together with the nine proteins shared by siKRAS_ and siCTRL_ctrlCM cells, though with opposite expression tendencies, revealed that the upregulated proteins are mainly localized at the mitochondrial matrix, whereas the downregulated ones were heterogeneously distributed by cell division structures (microtubule, centralspindlin and kinesin complexes and the kinetochore), as well as at the mitochondrion and mitochondrial matrix, among others. Concerning the biological process category, upregulated proteins were involved in mitotic cell cycle checkpoint, response to amino acid stimuli, gamma-aminobutyric acid biosynthesis, negative regulation of catalytic activity, mitotic DNA replication checkpoint and negative regulation of DNA replication. Owing to their location, downregulated proteins were associated with many biological processes such as, mitotic cytokinesis, microtubule-based movement, actomyosin contractile ring assembly, cell division, cytokinesis, mitotic nuclear division and mitotic spindle assembly, negative regulation of apoptotic processes, retrograde vesicle-mediated transport Golgi to endoplasmic reticulum and others. As for the molecular function, only downregulated proteins were significantly involved in microtubule, beta-tubulin, protein, poly(A) RNA, neuregulin and chaperone binding, as well as ATPase and microtubule motor activity. KEGG pathways analysis showed that upregulated proteins were involved in lysosome, while the downregulated ones were involved in ECM-receptor interaction pathways (Figure 6d).

Moreover, the 100 exclusive proteins of siKRAS_FibCM cells together with the nine common proteins with opposite expression tendencies, were mainly localized at the cytosol, extracellular exosomes, and focal adhesions (upregulated proteins); whilst downregulated proteins displayed more disperse locations at transport vesicles, cell-cell junctions, the extrinsic compartment of the cytoplasmic site of the plasma membrane, focal adhesions, nucleolus and at the small nucleolar ribonucleoprotein complexes. Upregulated proteins were mainly involved on synaptomenal complex disassembly, positive regulation of gene expression and transcription factors import to the nucleus whereas actin filament polymerization was a biologic process mainly observed in downregulated proteins. Concerning molecular function category, upregulated proteins were involved in protein, zinc, identical protein and ATP binding and aldehyde dehydrogenase activity, while downregulated proteins only showed significative results regarding poly(A) RNA binding. KEGG pathways analysis showed that upregulated proteins were involved in proteoglycans in cancer and p53 signaling pathway, while the downregulated proteins were involved in Rap1 signaling pathway (Figure 6e). Overall, these results show that KRAS-silenced cells respond to fibroblast-secretome by modulating transcription and by decreasing the expression of proteins associated with the cytoskeleton, focal adhesions and cell-cell junctions, thus representing key proteins involved in essential cancer cell functions, as motility.

The abovementioned analyses and rational was also followed to study the effect of fibroblast-secretome on the modulation of KRAS oncogenic proteome in LS174T cell line. The comparison of siKRAS vs siCTRL in ctrlCM, pinpointed 84 significatively altered proteins (43 upregulated and 41 downregulated). The impact of FibCM resulted in 63 significatively deregulated proteins (33 upregulated and 30 downregulated) between siKRAS and siCTRL cells (Figure 7a and Supplementary File S2). The comparative analysis of these two datasets showed that 63 proteins were exclusively regulated by KRAS in ctrlCM and 42 were exclusively regulated by KRAS in response to FibCM (Figure 7b). Twenty-one proteins were common to both conditions, from which 14 followed the same tendency and seven followed opposite expression tendencies. Inclusion of these seven proteins in the group of proteins exclusively found in siKRAS_FibCM cells revealed that 78% (49/63) of KRAS-associated proteome is fibroblast-secreted factors dependent (Figure 7a-e and Supplementary File S2). Five out of seven proteins with opposite tendencies, were downregulated upon culture with FibCM (ALDOB, MFN1, TNFRSF10B, TRMU and MFF) and significantly associated with mitochondrial outer membrane and involved in mitochondrial fusion (Figure 7b). The 14 shared proteins that follow the same tendency (ZMYM3, VPS37, RNASE1, UGT2B17, DEFA6, GDE1, B3GLCT, KDELC2, PCDH1, SRPK2, DNAJB4, QSER1, KIF1C and FUOM) did not show any significant association (Figure 7c).

**Figure 7:**
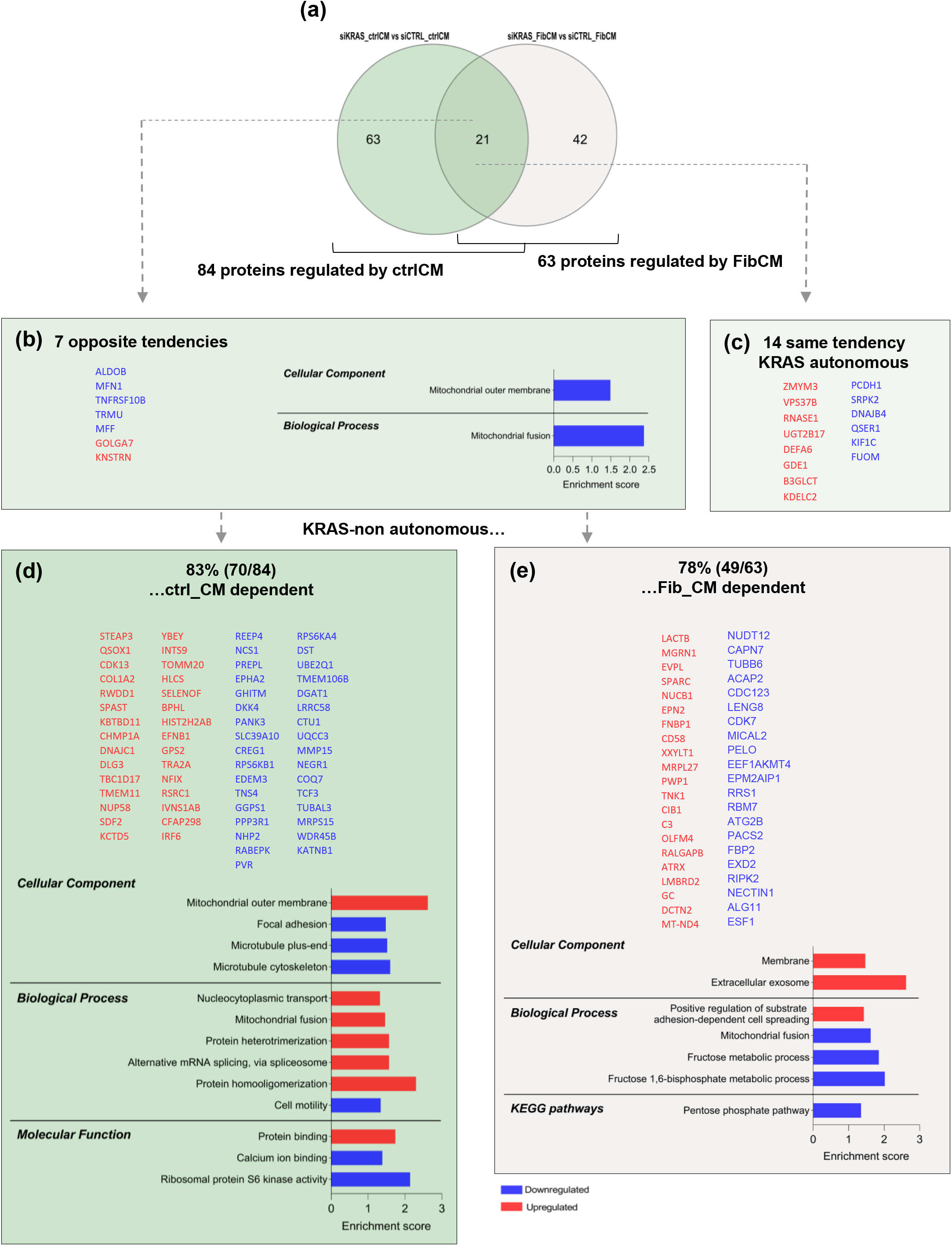
Analysis of differentially expressed proteins in LS174T cells, when comparing siKRAS_ctrlCM vs siCTRL_ctrlCM and siKRAS_FibCM vs siCTRL_FibCM. (a) Venn diagram showing exclusive and common proteins identified in the analyzed conditions. (b-e) GO and KEGG pathways analysis of: common proteins following opposite trends (b), common proteins following the same tendency, showing no significant results (c), exclusive proteins together with the commons that followed opposite tendencies found in the comparison of siKRAS vs siCTRL cultured in ctrlCM (d) and exclusive proteins together with the commons that followed opposite tendencies found in the comparison of siKRAS vs siCTRL cultured in FibCM (e). The enrichment score was calculated by –log P-value.

Analyses of the 63 siKRAS_ctrlCM exclusive proteins together with the seven proteins with opposite tendencies, showed that upregulated proteins localize at the mitochondrial outer membrane, while the downregulated are at the microtubule cytoskeleton and plus-end and at the focal adhesions. Upregulated proteins were further found to be related with protein homoligomerization and heterotrimerization, alternative mRNA splicing, mitochondrial fusion and nucleocytoplasmic transport; whilst downregulated proteins were involved in cell motility. Regarding the molecular function, upregulated proteins were shown to be associated with protein binding, and the downregulated ones with ribosomal protein S6 Kinase activity and calcium ion binding (Figure 7d). GO analyses of the 42 exclusive proteins of siKRAS_FibCM cells and the seven shared proteins with opposite tendencies were only statistically significant deregulated in cellular component and biological process categories. For cellular components, only upregulated proteins showed significance, being localized at extracellular exosomes and at the membrane. As for biological processes, upregulated proteins were mainly involved in positive regulation of substrate adhesion-dependent cell-spreading, while downregulated proteins were involved in fructose metabolism and mitochondrial fusion. In accordance, KEGG pathways analysis showed that downregulated proteins belong to the pentose phosphate pathway (Figure 7e). Hence, LS174T siKRAS cells likely respond to the fibroblast-secreted factors by adapting their metabolism.

Overall, our results suggest that, in both cell lines, the proteome associated to oncogenic KRAS is mainly regulated in a KRAS non-autonomous manner, as the expression of most proteins was found to be dependent on fibroblast-derived signals. Moreover, we observed that both KRAS silencing, and fibroblast-secreted factors had more impact on the proteome of HCT116 cells, as more proteins were differentially expressed in comparison with LS174T cells. All together, these results reinforce the complexity and the high intrinsic and extrinsic context-dependency of mutKRAS-driven effects.

## Discussion

While cancer cells KRAS-autonomous signaling has been extensively studied, KRAS non-autonomous effects, meaning how mutKRAS cancer cells respond to extrinsic factors, are less approached. The study of the interplay between mutKRAS cancer cells and the components of the TME is worth to the understanding of KRAS-driven non-autonomous signaling and represents a valuable strategy to understand therapy responses and to uncover putative therapeutical targets for mutKRAS cancer patients. Herein, we investigated the influence of TGFβ1-activated fibroblast-secretome on the proteome of mutKRAS CRC cell lines, thus addressing the context dependency of KRAS-associated signaling. Our data shows that the proteome alterations observed upon KRAS-silencing, depend not only on the cell line studied, but also on the external factors that cells are exposed to, highlighting the relevance of microenvironmental cues in KRAS-associated effects. In this study we resorted to a normal-like colon fibroblast cell line (CCD-18Co) and to the activation with TGFβ1 to mimic CAFs. In fact, TGFβ1 is the main driver of CAFs-associated features (e.g.: increased production of ECM components, expression of the α-SMA with and associated contraction capacity) [18]; however, one can under-stand that despite the biological relevance of this model, it is not representative of the heterogeneity of CAFs populations. Despite representing a limitation of this study, it dissolves the variability associated with the use of primary CAFs isolated from tumor patients with different genetic backgrounds, which can translate into significative dif-ferences in CAFs “pre-education”, resulting in conflicting results.

Our data shows that independently of direct cell-cell contact, TGFβ1-activated fibroblasts secretome was able to differently modulate the proteome of two CRC cell lines. Several aspects may explain their distinct responses. For instance, the type of KRAS mutation: G13D in HCT116 and G12D in LS174T [19]. These mutations present biochemical and structural differences; while the G12D has low affinity to RAF and a fast hydrolysis rate, the G13D has high affinity to RAF and a more rapid GTPase activity [6]. Also, drug sensitivity and therapy response of these mutant forms have been shown to be different [20–22]. In addition, these cell lines belong to different consensus molecular subtypes (CMS)-HCT116 belongs to the CMS4, the mesenchymal type, known to be enriched in fibroblasts, while LS174T is classified as CMS3, the metabolic type [23]. In accordance, HCT116 (both siCTRL and siKRAS) were more affected by the fibroblast-derived factors; whilst in LS174T, many of the alterations found, were related with metabolic pathways. Interestingly, fibroblast-secreted factors underlined the upregulation of RAF-1 in HCT116 cells supporting the role of KRAS downstream signaling pathways in the response to microenvironmental factors. Moreover, in both cell lines, proteins related to ECM (COL1A1, FN1 and TGFBI) were found upregulated. Individually, LS174T also showed upregulation of COL1A2, MMP2, DCN, TIMP-1 and other proteins related to ECM production/remodeling. For instance, the TGFβ1-driven ECM protein-TGFBI, found upregulated in both cell lines, has been shown to play important roles in essential processes underlying CRC metastasis formation, such as angiogenesis [24] and extravasation [25]. In line with these effects, TGBI expression is an independent poor prognostic factor in CRC [26]. As for TIMP-1, found in LS174T cells, it was already shown to promote tumor progression in prostate and colon cancer models, by driving the accumulation of CAFs [27]. So, one can infer that activated-fibroblasts secretome is capable to educate cancer cells towards tumor progression-related behaviors.

By addressing the influence of mutKRAS on the modulation of the proteome responsiveness to fibroblast-secreted factors, our data showed that almost 43% and 37,7% of the proteins modulated by fibroblast-secreted factors in HCT116 and LS174T cells, respectively, is indeed dependent on the presence of oncogenic KRAS. Given the important role of the tumor microenvironment derived signals to drive tumor malignant features [28], these results indicate that KRAS targeted inhibition can partially abrogate the protumorigenic stimuli derived from the tumor microenvironment. Still, more than a half of the proteome was found to be modulated by external factors in a KRAS-independent manner. Therefore, it is important to determine whether these KRAS-independent modifications contribute to induce resistance to KRAS inhibition. For example, in HCT116, FibCM led to the upregulation of proteins involved in cellular response to epidermal growth factor stimulus and substrate adhesion-dependent cell spreading. In LS174T, upregulated proteins were involved in several processes like extracellular matrix organization and disassembly, cell adhesion, wound healing, angiogenesis and blood vessel development, collagen catabolic process and fibril organization. These data indicate that FibCM upregulates pathways that may endow KRAS-inhibited tolerant cancer cells with pro-malignant features. Our data shows that, upon KRAS silencing and exposure to FibCM, the responses of the two cell lines seem, in some aspects, to go in opposite directions: HCT116 cells show a significant downregulation of Rap-1 signaling (an important signaling pathway in cancer malignancy, involved for instance in epithelial to mesenchymal transition, invasion and angiogenesis promotion [29]); LS174T cells upregulate proteins that are involved in adhesion and positive cell spreading regulation, counteracting the downregulation of proteins involved in cell motility when siKRAS cells are cultured in ctrCM. On top of reinforcing the context dependency of KRAS signaling, these results show that in some contexts, fibroblast-derived external stimuli can, in fact, modulate the cancer cell proteome in a way that counteract KRAS-inhibition and support cancer cell-malignant features. This goes in line with our previous results that demonstrated opposite invasive responses of HCT116 and LS174T to fibroblast-secreted factors upon KRAS silencing. While in HCT116, KRAS inhibition impaired invasion induced by fibroblasts-derived HGF, in LS174T KRAS inhibition triggered invasion [16]. Moreover, both cell lines commonly upregulated proteins associated with exosomes localization upon silencing of KRAS and culture with FibCM. Since exosomes play an important role in cell-to-cell communication both locally and at distance (e.g. in metastatic niche preparation) [30–32], it is worth to profile the content of the exosomes produced in this context.

In addition, our data show that about 2/3 of the proteome associated to oncogenic KRAS is controlled by external factors. These results suggest that, in tumors, oncogenic KRAS signaling may be very heterogeneous as, within the tumor bulk, cells may engage different oncogenic KRAS signaling programs according to the microenvironmental niche they are exposed to. For instance, this context-dependent heterogeneity may dictate different responses to therapy. From the analysis on HCT116 KRAS silenced cells, one of the upregulated pathways highlighted as KRAS-autonomous, is related with type I interferon signaling. Importantly, in CRC models, an interferon gene expression signature was shown to underlie MEK inhibition resistance in a mutKRAS context [33]. As such, together with our result, this suggests a KRAS pathway inhibition resistance mechanism, thus contributing to explain the failure of KRAS pathway inhibitors and suggesting a possible combinatorial treatment. Moreover, this response was cell-line specific, as it was not observed in LS174T cells. Also, in HCT116 cells, one protein that stands out from the common proteins found upon KRAS silencing independently of FibCM following opposite directions is the Polo Like Kinase 1 (or Serine/threonine-protein kinase PLK1). Upon KRAS silencing PLK1 expression was downregulated, however the exposure to FibCM rescues that effect causing an upregulation of this kinase. MutKRAS cancer cells have been shown to be sensitive to PLK1 inhibition [34,35]. Hence, the upregulation induced by the exposure to fibroblast-secreted factors, might constitute an evasion mechanism to KRAS signaling inhibition. Indeed, a phase 1b/2clinical trial (NCT03829410) is currently evaluating the safety and efficacy of the PLK1 inhibitor onvansertib in combination with FOLFIRI + Bevacizumab, in mutKRAS metastatic CRC. Notwithstanding, additional studies are needed to determine how mutant KRAS cells respond to other microenvironment cell types and how it affects the response to KRAS direct targeting or inhibition of its downstream effectors. More so, the identification of a KRAS-autonomous signature common to different types of stimuli is likely to reveal valuable actionable targets that can be used to impair mutant KRAS cells irrespective of their signaling heterogeneity induced by different microenvironments.

Gathering our data, one can appreciate that most of the proteomic profile alterations resulted from KRAS non-autonomous signaling, reinforcing the importance of re-evaluating oncogenic KRAS signaling in the context of the tumor microenvironment. This will certainly improve the understanding of mutKRAS CRC biology and empower the identification of therapeutic targets and disease biomarkers of therapy response, allowing a better patient stratification. Functional validation of the disclosed targets in future studies, as well as the study of the secretome components behind such effects, will provide insights on their therapeutical relevance, either as targets or biomarkers.

## Conclusions

In sum, this work constitutes a step forward towards the understanding of KRAS-associated signals in response to microenvironmental cues. Also, it highlights the need of the development of more complex models that can faithfully mimic the complexity of the TME to study the behavior of mutKRAS cancer cells and generate translational knowledge to be used in the identification of therapeutic targets or biomarkers of resistance to therapy.

## Acknowledgments

The authors acknowledge the i3S Proteomics Scientific Platform. The mass spectrometry technique was performed by Hugo Osório and the work had support from the Portuguese Mass Spectrometry Network, integrated in the National Roadmap of Research Infrastructures of Strategic Relevance (ROTEIRO/0028/2013; LISBOA-01-0145-FEDER-022125).

## SUPPLEMENTARY MATERIALS

Figure S1: Representative western blot showing increased expression of α-SMA following CCD-18Co fibroblasts activation with rhTGFβ1.

Figure S2: Representative western blots showing efficient KRAS silencing in HCT116 and LS174T cell lines.

Supplementary File S1: Lists of differentially expressed proteins in HCT116 cell line. Supplementary File

S2: Lists of differentially expressed proteins in LS174T cell line.

## References

1. Hobbs, G.A.; Der, C.J.; Rossman, K.L. RAS isoforms and mutations in cancer at a glance. J. Cell Sci. 2016, 129, 1287–1292, doi:10.1242/jcs.182873.

2. Haigis, K.M. KRAS Alleles: The Devil Is in the Detail. Trends in Cancer 2017, 3, 686–697, doi:10.1016/j.trecan.2017.08.006.

3. Moore, A.R.; Rosenberg, S.C.; McCormick, F.; Malek, S. RAS-targeted therapies: is the undruggable drugged? Nat. Rev. Drug Discov. 2020, 19, 533–552, doi:10.1038/s41573-020-0068-6.

4. Giehl, K.; Skripczynski, B.; Mansard, A.; Menke, A.; Gierschik, P. Growth factor-dependent activation of the Ras-Raf-MEK-MAPK pathway in the human pancreatic carcinoma cell line PANC-1 carrying activated K-ras: Implications for cell proliferation and cell migration. Oncogene 2000, 19, 2930–2942, doi:10.1038/sj.onc.1203612.

5. Smith, M.J.; Ikura, M. Integrated RAS signaling defined by parallel NMR detection of effectors and regulators. Nat. Chem. Biol. 2014, 10, 223–230, doi:10.1038/nchembio.1435.

6. Hunter, J.C.; Manandhar, A.; Carrasco, M.A.; Gurbani, D.; Gondi, S.; Westover, K.D. Biochemical and structural analysis of common cancer-associated KRAS mutations. Mol. Cancer Res. 2015, 13, 1325–1335, doi:10.1158/1541-7786.MCR-15-0203.

7. Yuan, T.L.; Amzallag, A.; Bagni, R.; Yi, M.; Afghani, S.; Burgan, W.; Fer, N.; Strathern, L.A.; Powell, K.; Smith, B.; et al. Differential Effector Engagement by Oncogenic KRAS. Cell Rep. 2018, 22, 1889–1902, doi:10.1016/j.celrep.2018.01.051.

8. Brubaker, D.K.; Paulo, J.A.; Sheth, S.; Poulin, E.J.; Popow, O.; Joughin, B.A.; Strasser, S.D.; Starchenko, A.; Gygi, S.P.; Lauffenburger, D.A.; et al. Proteogenomic Network Analysis of Context-Specific KRAS Signaling in Mouse-to-Human Cross-Species Translation. Cell Syst. 2019, 9, 258–270.e6, doi:10.1016/j.cels.2019.07.006.

9. Cook, J.H.; Melloni, G.E.M.; Gulhan, D.C.; Park, P.J.; Haigis, K.M. The origins and genetic interactions of KRAS mutations are allele-and tissue-specific. Nat. Commun. 2021, 12, doi:10.1038/s41467-021-22125-z.

10. Kim, E.; Kim, J.Y.; Smith, M.A.; Haura, E.B.; Anderson, A.R.A. Cell signaling heterogeneity is modulated by both cell-intrinsic and -extrinsic mechanisms: An integrated approach to understanding targeted therapy. PLoS Biol. 2018, 16, 1–29, doi:10.1371/journal.pbio.2002930.

11. Dias Carvalho, P.; Guimarães, C.F.; Cardoso, A.P.; Mendonça, S.; Costa, A.M.; Oliveira, M.J.; Velho, S. KRAS Oncogenic Signaling Extends beyond Cancer Cells to Orchestrate the Microenvironment. Cancer Res. 2018, 78, 7–14, doi:10.1158/0008-5472.CAN-17-2084.

12. Dias Carvalho, P.; Machado, A.L.; Martins, F.; Seruca, R.; Velho, S. Targeting the Tumor Microenvironment: An Unexplored Strategy for Mutant KRAS Tumors. Cancers (Basel). 2019, 11, 2010, doi:10.3390/cancers11122010.

13. Deng, S.; Clowers, M.J.; Velasco, W. V; Ramos-castaneda, M. Understanding the Complexity of the Tumor Microenvironment in K-ras Mutant Lung Cancer : Finding an Alternative Path to Prevention and Treatment. Front. Oncol. 2020, 9, 1–17, doi:10.3389/fonc.2019.01556.

14. Hamarsheh, S.; Groß, O.; Brummer, T.; Zeiser, R. Immune modulatory effects of oncogenic KRAS in cancer. Nat. Commun. 2020, 11, 5439, doi:10.1038/s41467-020-19288-6.

15. Liu, H.; Liang, Z.; Zhou, C.; Zeng, Z.; Wang, F.; Hu, T.; He, X.; Wu, X.; Wu, X.; Lan, P. Mutant KRAS triggers functional reprogramming of tumor-associated macrophages in colorectal cancer. Signal Transduct. Target. Ther. 2021, 6, 1–13, doi:10.1038/s41392-021-00534-2.

16. Dias Carvalho, P.; Martins, F.; Mendonça, S.; Ribeiro, A.; Machado, A.L.; Carvalho, J.; Oliveira, M.J.; Velho, S. Mutant KRAS modulates colorectal cancer cells invasive response to fibroblast-secreted factors through the HGF/C-MET axis. bioRxiv 2021, 2021.11.16.468815, doi:10.1101/2021.11.16.468815.

17. Perez-Riverol, Y.; Csordas, A.; Bai, J.; Bernal-Llinares, M.; Hewapathirana, S.; Kundu, D.J.; Inuganti, A.; Griss, J.; Mayer, G.; Eisenacher, M.; et al. The PRIDE database and related tools and resources in 2019: Improving support for quantification data. Nucleic Acids Res. 2019, 47, D442–D450, doi:10.1093/nar/gky1106.

18. Denys, H.; Derycke, L.; Hendrix, A.; Westbroek, W.; Gheldof, A.; Narine, K.; Pauwels, P.; Gespach, C.; Bracke, M.; De Wever, O. Differential impact of TGF-B1 and EGF on fibroblast differentiation and invasion reciprocally promotes colon cancer cell invasion. Cancer Lett. 2008, 266, 263–274, doi:10.1016/j.canlet.2008.02.068.

19. Ahmed, D.; Eide, P.W.; Eilertsen, I.A.; Danielsen, S.A.; Eknæs, M.; Hektoen, M.; Lind, G.E.; Lothe, R.A. Epigenetic and genetic features of 24 colon cancer cell lines. Oncogenesis 2013, 2, 1–8, doi:10.1038/oncsis.2013.35.

20. Garassino, M.C.; Marabese, M.; Rusconi, P.; Rulli, E.; Martelli, O.; Farina, G.; Scanni, A.; Broggini, M. Different types of K-Ras mutations could affect drug sensitivity and tumour behaviour in non-small-cell lung cancer. Ann. Oncol. 2011, 22, 235–237, doi:doi:10.1093/annonc/mdq680.

21. Duldulao, M.P.; Lee, W.; Nelson, R.A.; Li, W.; Chen, Z.; Kim, J.; Garcia-Aguilar, J. Mutations in specific codons of the KRAS oncogene are associated with variable resistance to neoadjuvant chemoradiation therapy in patients with rectal adenocarcinoma. Ann. Surg. Oncol. 2013, 20, 2166–2171, doi:doi:10.1245/s10434-013-2910-0.

22. Mao, C.; Huang, Y.F.; Yang, Z.Y.; Zheng, D.Y.; Chen, J.Z.; Tang, J.L. KRAS p.G13D mutation and codon 12 mutations are not created equal in predicting clinical outcomes of cetuximab in metastatic colorectal cancer: A systematic review and meta-analysis. Cancer 2013, 119, 714–721, doi:10.1002/cncr.27804.

23. Berg, K.C.G.; Eide, P.W.; Eilertsen, I.A.; Johannessen, B.; Bruun, J.; Danielsen, S.A.; Bjørnslett, M.; Meza-Zepeda, L.A.; Eknæs, M.; Lind, G.E.; et al. Multi-omics of 34 colorectal cancer cell lines - a resource for biomedical studies. Mol. Cancer 2017, 16, 1–16, doi:10.1186/s12943-017-0691-y.

24. Chiavarina, B.; Costanza, B.; Ronca, R.; Blomme, A.; Rezzola, S.; Chiodelli, P.; Giguelay, A.; Belthier, G.; Doumont, G.; van Simaeys, G.; et al. Metastatic colorectal cancer cells maintain the TGFβ program and use TGFBI to fuel angiogenesis. Theranostics 2021, 11, 1626–1640, doi:10.7150/thno.51507.

25. Ma, C.; Rong, Y.; Radiloff, D.R.; Datto, M.B.; Centeno, B.; Bao, S.; Cheng, A.W.M.; Lin, F.; Jiang, S.; Yeatman, T.J.; et al. Extracellular matrix protein βig-h3/TGFBI promotes metastasis of colon cancer by enhancing cell extravasation. Genes Dev. 2008, 22, 308–321, doi:10.1101/gad.1632008.

26. Zhu, J.; Chen, X.; Liao, Z.; He, C.; Hu, X. TGFBI protein high expression predicts poor prognosis in colorectal cancer patients. Int. J. Clin. Exp. Pathol. 2015, 8, 702–710.

27. Gong, Y.; Scott, E.; Lu, R.; Xu, Y.; Oh, W.K.; Yu, Q. TIMP-1 Promotes Accumulation of Cancer Associated Fibroblasts and Cancer Progression. PLoS One 2013, 8, e77366, doi:10.1371/journal.pone.0077366.

28. Hanahan, D.; Coussens, L.M. Accessories to the Crime: Functions of Cells Recruited to the Tumor Microenvironment. Cancer Cell 2012, 21, 309–322, doi:10.1016/j.ccr.2012.02.022.

29. Looi, C.; Hii, L.; Ngai, S.C.; Leong, C.; Mai, C. The Role of Ras-Associated Protein 1 (Rap1) in Cancer: Bad Actor or Good Player? Biomedicines 2020, 8, 334, doi:10.3390/biomedicines8090334.

30. Beckler, M.D.; Higginbotham, J.N.; Franklin, J.L.; Ham, A.J.; Halvey, P.J.; Imasuen, I.E.; Whitwell, C.; Li, M.; Liebler, D.C.; Coffey, R.J. Proteomic analysis of exosomes from mutant KRAS colon cancer cells identifies intercellular transfer of mutant KRAS. Mol. Cell. Proteomics 2013, 12, 343–355, doi:10.1074/mcp.M112.022806.

31. Hoshino, A.; Costa-Silva, B.; Shen, T.-L.; Rodrigues, G.; Hashimoto, A.; Mark, M.T.; Molina, H.; Kohsaka, S.; Giannatale, A. Di; Ceder, S.; et al. Tumour exosome integrins determine organotropic metastasis. Nature 2015, 527, 329–335, doi:10.1038/nature15756.Tumour.

32. Shao, Y.; Chen, T.; Zheng, X.; Yang, S.; Xu, K.; Chen, X.; Xu, F.; Wang, L.; Shen, Y.; Wang, T.; et al. Colorectal cancer-derived small extracellular vesicles establish an inflammatory premetastatic niche in liver metastasis. Carcinogenesis 2018, 39, 1368–1379, doi:10.1093/carcin/bgy115.

33. Wagner, S.; Vlachogiannis, G.; Haven, A. De; Melanie, B.; Gary, V.; Jenkins, L.; Mancusi, C.; Self, A.; Manodoro, F.; Assiotis, I.; et al. Suppression of interferon gene expression overcomes resistance to MEK inhibition in KRAS -mutant colorectal cancer. Oncogene 2019, 38, 1717–1733, doi:10.1038/s41388-018-0554-z.

34. Luo, J.; Emanuele, M.J.; Li, D.; Creighton, C.J.; Schlabach, M.R.; Westbrook, T.F.; Wong, K.K.; Elledge, S.J. A Genome-wide RNAi Screen Identifies Multiple Synthetic Lethal Interactions with the Ras Oncogene. Cell 2009, 137, 835–848, doi:10.1016/j.cell.2009.05.006.

35. Wang, J.; Hu, K.; Guo, J.; Cheng, F.; Lv, J.; Jiang, W.; Lu, W.; Liu, J.; Pang, X.; Liu, M. Suppression of KRas-mutant cancer through the combined inhibition of KRAS with PLK1 and ROCK. Nat. Commun. 2016, 7, 11363, doi:10.1038/ncomms11363.

