## Supplementary figures for "Mutant KRAS-associated proteome is mainly controlled by exogenous factors"

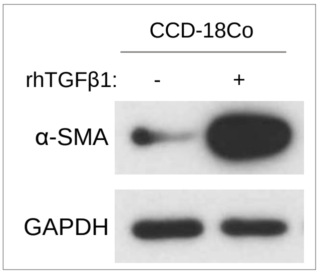


**Figure S1:** Representative western blot showing increased expression of α-SMA following CCD-18Co fibroblasts activation with rhTGFβ1.

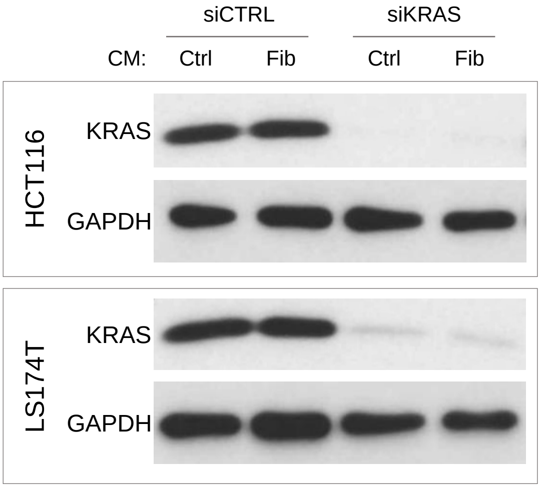


**Figure S2:** Representative western blots showing efficient KRAS silencing in HCT116 and LS174T cell lines.
